# BRCA1/BARD1 ubiquitinates PCNA in unperturbed conditions to promote replication fork stability and continuous DNA synthesis

**DOI:** 10.1101/2023.01.12.523782

**Authors:** Daniel Salas-Lloret, Néstor García-Rodríguez, Lisanne Giebel, Arnoud de Ru, Peter A. van Veelen, Pablo Huertas, Alfred C.O. Vertegaal, Román González-Prieto

## Abstract

Deficiencies in the BRCA1 tumor suppressor gene are the main cause of hereditary breast and ovarian cancer. BRCA1 is involved in the Homologous Recombination DNA repair pathway, and, together with BARD1, forms a heterodimer with ubiquitin E3 activity. The relevance of the BRCA1/BARD1 ubiquitin E3 activity for tumor suppression and DNA repair remains controversial and most efforts aimed to identify BRCA1/BARD1 ubiquitination substrates rely on indirect evidence. Here, we observed that the BRCA1/BARD1 ubiquitin E3 activity was not required for Homologous Recombination or resistance to Olaparib. Using TULIP2 methodology, which enables the direct identification of E3-specific ubiquitination substrates, we identified substrates for BRCA1/BARD1. We found that PCNA is ubiquitinated by BRCA1/BARD1 in unperturbed conditions independently of RAD18. PCNA ubiquitination by BRCA1/BARD1 avoids the formation of ssDNA gaps during DNA replication and promotes replication fork stability epistatically to BRCA1 S114 phosphorylation, addressing the controversy about the function of BRCA1/BARD1 E3 activity in Homologous Recombination.

## INTRODUCTION

Breast cancer susceptibility type 1 (BRCA1) binds its partner BRCA1-Associated RING Domain 1 (BARD1) to form an obligated and multifunctional heterodimer with ubiquitin E3 ligase activity (Brzovic et al., 2001; Densham et al., 2016; Noordermeer et al., 2018). The BRCA1/BARD1 heterodimer acts as tumor suppressor and maintains genome stability, generally, by repairing deleterious double-strand DNA breaks (DSBs) via error-free homologous recombination (HR) (Tarsounas and Sung, 2020; Zhao et al., 2017). DSBs can originate either from endogenous agents, such as reactive oxygens, replication fork progression and single-stranded DNA (ssDNA), or from environmental exposure to chemicals, ionizing radiation and ultraviolet light (UV) (Tubbs and Nussenzweig, 2017).

Germline mutations in BRCA1 and BARD1 are the main cause of hereditary breast and ovarian cancers, conferring a life-time probability of up to 90% for developing breast cancer and 50% for ovarian cancer (King et al., 2003; Scully and Puget, 2002; Weber-Lassalle et al., 2019). Mice studies suggested that the E3 ligase activity is not essential to prevent tumor development (Shakya et al., 2011). However, mutations in BARD1 that impair Histone 2A (H2A) ubiquitination have been identified in familial breast cancer (Stewart et al., 2018).

BRCA1/BARD1 has a very well established histone H2A ubiquitin ligase activity on lysines K127/K129 (Kalb et al., 2014; Kim et al., 2017; Witus et al., 2021; Zhu et al., 2011). This ubiquitination has been related to the maintenance of heterochromatin integrity, genetic stability and senescence (Densham *et al*., 2016; Kim *et al*., 2017; Zhu *et al*., 2011).

Up to date, the relevance of the BRCA1/BARD1 E3 activity for DNA Double Strands Break repair, tumor suppression and resistance to PARP inhibitors and platinum-based compounds is still controversial. Histone H2A K127/129 ubiquitination is required for DNA DSB repair by homologous recombination and RAD51 foci formation (Densham *et al*., 2016). However, deficiencies in histone H2A ubiquitination by BRCA1/BARD1 can be rescued by other ubiquitin E3s such as RNF168 (Sherker et al., 2021; Zong et al., 2019). While a study from the Morris group (Densham *et al*., 2016) using siRNA-based knockdowns in HeLa cells showed that the BRCA1/BARD1 E3 activity was required for resistance to agents such as the PARP inhibitor Olaparib and cisplatin, a degrON-based strategy on HCT116 cells showed that the BRCA1/BARD1 E3 activity was dispensable for resistance to Olaparib (Nakamura et al., 2019), consistent with most of the published literature (Drost et al., 2011; Nakamura *et al*., 2019; Reid et al., 2008; Shakya *et al*., 2011; Sherker *et al*., 2021).

Moreover, many research groups have addressed the challenge of identifying BRCA1/BARD1 ubiquitination substrates. However, all these studies have relied in indirect evidence to identify putative BRCA1/BARD1 ubiquitination substrates, by either overexpressing or depleting BRCA1 and identification of changes in the ubiquitin proteome by mass spectrometry-based proteomics.(Barrows et al., 2021; Eakin et al., 2007; Heidelberger et al., 2016; Kim *et al*., 2017; Matsuzawa et al., 2014; Song et al., 2011). However, since BRCA1/BARD1 plays an important role in several signaling pathways including cell cycle regulation, replication fork protection, gene transcription regulation and DNA damage repair (Deng, 2006; Tarsounas and Sung, 2020), these approaches could indirectly affect the ubiquitination state of proteins. Recently, we developed the TULIP and TULIP2 methodologies (Kumar et al., 2017; Salas-Lloret et al., 2019) which enabled the unambiguous identification of direct ubiquitination substrates for an E3 of interest.

Here, we investigated the role of the BRCA1/BARD1 heterodimer E3 activity in the non-tumoral cell line RPE1 and applied the TULIP2 methodology to identify novel substrates for ubiquitination by BRCA1/BARD1. Combined, our data provided novel insights in BRCA1/BARD1 function and enabled us to solve the controversy in the field about the relevance of the ubiquitin E3 function of BRCA1/BARD1 for homologous recombination.

## RESULTS

### BRCA1/BARD1 ubiquitin E3 activity is dispensable for DSB repair and resistance to treatment with Olaparib

The relevance of the BRCA1/BARD1 ubiquitin E3 activity for homologous recombination in terms of DNA DSB repair and treatment resistance remains contradictory in current literature applying different approaches in different cancer cell lines and models (Densham *et al*., 2016; Drost *et al*., 2011; Nakamura *et al*., 2019; Reid *et al*., 2008; Shakya *et al*., 2011). To address this controversy, we employed RPE1 *TP53^-/-^* (Parental) and RPE1 *TP53^-/-^ BRCA1^-/-^* (BRCA1-KO) cells (Noordermeer *et al*., 2018) rescued or not with GFP-tagged BRCA1 wild type or I26A mutant. The BRCA1 I26A mutant abrogates the BRCA1/BARD1 E3 activity without disrupting the formation of the BRCA1/BARD1 heterodimer (Brzovic et al., 2003; Christensen et al., 2007). We decided to test the proficiency in homologous recombination of the ubiquitin E3 activity BRCA1 mutant I26A in non-tumoral cells both in terms of resistance to Olaparib (Figure 1A) and RAD51 foci formation in response to Ionizing Irradiation (IR) (Figure 1B-C). As expected, BRCA1-KO cells were deficient in resistance to Olaparib and RAD51 foci formation in response to ionizing radiation, and BRCA1-GFP rescued cells could restore both phenotypes to the Parental levels. Importantly, BRCA1_I26A_-GFP rescued cells could also restore Olaparib resistance and RAD51 foci to the Parental levels, which is consistent with most of the published literature about the relevance of BRCA1 ubiquitin E3 activity in homologous recombination in DSB repair and resistance to Olaparib and Platinum-based compounds (Drost *et al*., 2011; Nakamura *et al*., 2019; Reid *et al*., 2008; Shakya *et al*., 2011; Sherker *et al*., 2021).

**Figure 1.**
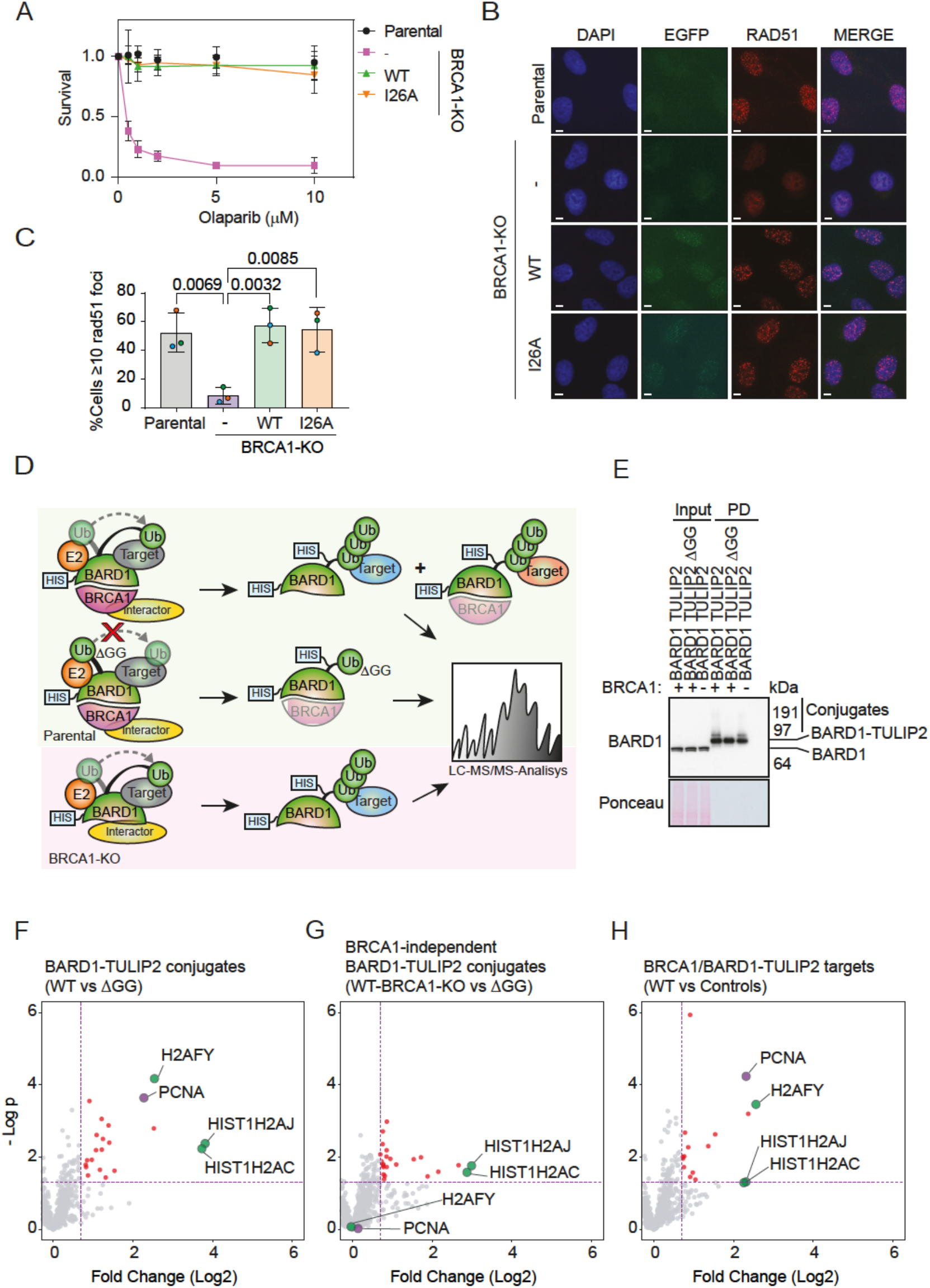
BRCA1 E3 activity is not required for homologous recombination. A. Clonogenic survival assay of Parental and BRCA1-KO cells complemented or not with either BRCA1-WT-GFP or BRCA1-I26A-GFP in response to different concentrations of Olaparib. Average and standard deviation of three independent experiments with three technical repeats are depicted (N=3). B-C. Immunofluorescence analysis Parental and BRCA1-KO cells complemented or nor with either BRCA1-WT-GFP or BRCA1-I26A-GFP after exposure to 10 Gy ionizing irradiation. Representative images (B) and quantification (C) of cells with RAD51 foci is shown. Size bars in fluorescence microscopy images represent 10 μm. Data corresponds to three independent experiments. The average of each individual experiment is shown by an orange, green or blue circle respectively. Unpaired t-tests were performed with p-values shown in the figure D. Cartoon depicting BARD1-TULIP2 rationale. E. Analysis by immunoblotting of BARD1-TULIP2 samples. F-H. Volcano plots depicting differences between each of the specified BARD1-TULIP2 constructs. Histones H2A and macro-H2A and PCNA are labelled. Each dot represents a protein.

### BRCA1/BARD1 ubiquitinates PCNA in a constitutive manner

Despite proficiency in DNA DSB repair, cancer predisposition has been observed in BRCA1 E3 mutant mouse models (Drost *et al*., 2011), and previous studies aiming to identify BRCA1/BARD1 ubiquitination substrates are based on indirect evidence (Kalb *et al*., 2014; Kim *et al*., 2017; Yu et al., 2006). Moreover, the ubiquitination of histone H2A-K127/K129, which is the best studied BRCA1/BARD1 ubiquitination substrate (Kalb *et al*., 2014; Kim *et al*., 2017), can be rescued by RNF168, another E3 enzyme, upon BRCA1 deficiencies (Zong *et al*., 2019). Thus, we hypothesized that BRCA1/BARD1 ubiquitination substrates other than histone H2A isoforms exist.

We recently developed the TULIP2 methodology for the systematic application of Ubiquitin Activated Interaction Traps (Kumar *et al*., 2017; O’Connor et al., 2015; Salas-Lloret *et al*., 2019), which enables the identification of E3-specific targets in a direct manner (Supplementary Figure 1A; Figure 1D). The rationale behind this approach is that if we have a linear fusion between an E3 and ubiquitin, this E3 could use its attached ubiquitin to modify its substrate, enabling the co-purification and subsequent identification by mass spectrometry-based proteomics of the E3 together with ubiquitin and the covalently modified substrate. Therefore, we decided to apply TULIP2 methodology to identify BRCA1/BARD1-specific ubiquitination substrates. Three different TULIP2 constructs were generated to identify BRCA1 direct ubiquitination targets: BRCA1-WT-TULIP2, BRCA1-I26A-TULIP2 and BRCA1-WT-TULIP2ΔGG. Both BRCA1-I26A and BRCA1-WT-TULIP2ΔGG served as negative controls. While BRCA1-I26A is a catalytic dead mutant without E3 activity, BRCA1-WT-TULIP2ΔGG fused ubiquitin cannot be conjugated to target proteins due to lack of the C-terminal diGly motif (Supplementary Figure 1A,B).

First, we confirmed the functionality of the BRCA1-TULIP2 constructs for homologous recombination by introducing them in BRCA1-KO cells in a stable inducible manner and testing their resistance to Olaparib treatment (Supplementary Figure 1C), and RAD51 foci formation in response to IR (Supplementary Figure 1D,E). As previously seen for the GFP-tagged constructs, BRCA1-KO cells were hypersensitive to Olaparib and failed to form RAD51 foci in response to IR. However, BRCA1-KO cells stably expressing either BRCA1-WT-TULIP2 or BRCA1-I26A-TULIP2 completely rescued both Olaparib sensitivity and deficiency in RAD51 foci formation (Supplementary Figure 1C-E). Besides, BRCA1-TULIP2 constructs co-localized with RAD51 similarly to endogenous BRCA1 (Supplementary Figure 1D). These results corroborate the functionality of BRCA1-TULIP2 constructs and indicate a minor role of BRCA1/BARD1 E3 activity on HR pathway for DNA DSB repair and Olaparib resistance.

Next, we performed the TULIP2 assay to identify BRCA1 ubiquitin E3 activity-specific substrates using our BRCA1-TULIP2 cell lines (Supplementary Figure 1A) and performed mass spectrometry-based proteomics analysis (Supplementary Dataset 1; Supplementary Figure 1F-G). Analysis by immunoblotting failed to show a smear up from the wild type and I26A corresponding to BRCA1-TULIP2 conjugates as expected from functional TULIPs (Supplementary Figure 1B) (González-Prieto and Vertegaal, 2019; Kumar *et al*., 2017; Salas-Lloret *et al*., 2019). Moreover, proteomics analysis was unable to identify neither histone H2A nor macro-H2A as BRCA1-TULIP2 substrates, which would had served as positive controls, and the top hit was RAB43, which is not a nuclear protein. We concluded that the BRCA1-TULIP2 constructs were functional for BRCA1 activity but not regarding the TULIP2 assay, likely due to steric hindrance.

Nevertheless, a recent cryo-EM study has shown that it is BARD1, and not BRCA1, the heterodimer partner which positions the E2 enzyme and directs the ubiquitination of histone H2A (Witus *et al*., 2021). Therefore, we made stable inducible BARD1-TULIP2 constructs and introduced them in Parental cells, including the wild type and ΔGG TULIP2 constructs. As an additional negative control, in this case, we introduced wild type BARD1-TULIP2 constructs in BRCA1-KO cells (Figure 1D). Analysis by immunoblotting of the BARD1-TULIP2-expressing cells (Figure 1E) revealed that: 1) The endogenous BARD1 levels were similar in parental and BRCA1-KO cells; 2) the BARD1-TULIP2 construct levels were not detectable compared to endogenous BARD1 levels in the input samples, which is important to avoid overexpression artefacts; and 3) The BARD1-TULIP2 constructs were very efficiently enriched in the His-pulldown following TULIP2 methodology and a smear up from the wild type BARD1-TULIP2 construct corresponding to BARD1-TULIP2 conjugates, which was absent in the ΔGG control, could be detected. Mass spectrometry analysis of BARD1-TULIP2 samples (Supplementary Dataset 2; Figure 1F-H) identified both histones H2A and macro-H2A as BARD1 ubiquitination substrates, serving as internal positive controls. Interestingly, we could identify another top hit, PCNA, as a BRCA1/BARD1specific ubiquitination substrate using TULIP2 methodology (Figure 1H).

### BRCA1/BARD1 and RAD18 ubiquitinates PCNA-K164 in distinct signaling pathways

The proteomics spectra obtained in our BARD1-TULIP2 experiments did not enable the identification of the ubiquitin acceptor lysine in PCNA for BRCA1/BARD1. K164 is the main ubiquitin acceptor site in PCNA, and this ubiquitination promotes the recruitment of Trans-Lesion Synthesis (TLS) DNA polymerases (Garcia-Rodriguez et al., 2016). However, other acceptor lysines for ubiquitin have been identified in PCNA, which physiological relevance are unknown (Akimov et al., 2018). Thus, there was the possibility that the BRCA1/BARD1 heterodimer was ubiquitinating PCNA on a lysine other than K164. Therefore, we performed the BARD1-TULIP2 assay on RPE1 *TP53^-/-^ PCNA^K164R/K164R^* (PCNA-K164R) cells (Thakar et al., 2020) (Supplementary Dataset 3; Figure 2A). In this assay, we could identify histones H2A and macroH2A as BARD1 ubiquitination substrates. However, PCNA was lost as a BARD1-specific substrate, which indicates that BRCA1/BARD1 ubiquitinates PCNA on lysine K164.

**Figure 2.**
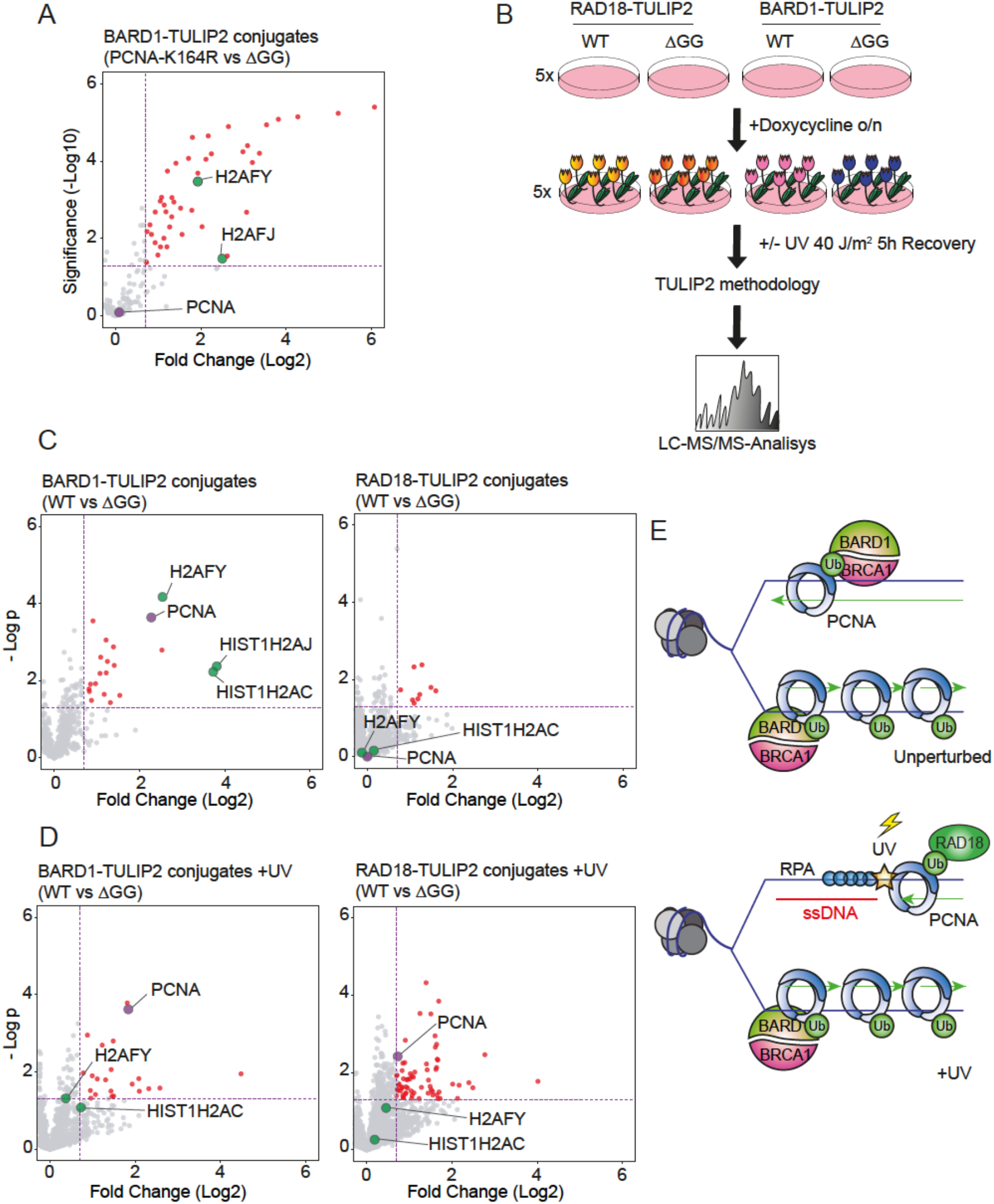
RAD18 and BARD1 ubiquitinates PCNA in different pathways. A. Volcano plot depicting differences between BARD1-TULIP2 conjugates in PCNA-K164R cell line versus DGG control. Histones H2A and macro-H2A and PCNA are labelled. B. Experimental setup. The expression of specified BARD1-TULIP2 and RAD18-TULIP2 constructs were induced overnight. Cells were irradiated with 40 J/m^2^ of UV light and allowed cells to recover for 5h prior to cell lysis, protein purification and identification by LC-MS/MS of the TULIP2 conjugates C, D. Volcano plots depicting differences between wild type BARD1-TULIP2 or RAD18-TULIP2 with their respective ΔGG counterparts in unperturbed conditions (C) or 5h after UV irradiation (D). Each dot represents a protein, PCNA, histone H2A and macro-H2A are labeled. E. Proposed model. During unperturbed conditions BRCA1/BARD1 ubiquitinates PCNA at low levels and upon UV irradiation RAD18 ubiquitinates PCNA in response to DNA lesions that represent a barrier for the replication fork while BRCA1/BARD1 still ubiquitinates PCNA.

BRCA1 has been shown to promote the recruitment of RAD18 to chromatin in response to replication-blocking lesions (Tian et al., 2013). Since RAD18 is considered the canonical ubiquitin ligase for PCNA in response to replication barriers (Choe and Moldovan, 2017; Lee and Myung, 2008; Watanabe et al., 2004) and protein-protein interactions between BRCA1 and RAD18 have been described (Tian *et al*., 2013), there was the possibility that it was RAD18 the E3 enzyme using the ubiquitin moiety in the BARD1-TULIP2 construct to modify PCNA. To exclude this possibility, we made RAD18-TULIP2 constructs, introduced them in Parental cells and performed the TULIP2 assay with and without UV light irradiation (Supplementary dataset 4; Figure 2B). As expected, immunoblotting analysis (Supplementary Figure 2A) showed that the wild type RAD18-TULIP2 samples formed a smear up from the RAD18-TULIP2 construct, indicative of its functionality, that was suppressed in the ΔGG negative controls. Mass spectrometry-based proteomics analysis of the non-irradiated samples revealed that among the identified RAD18 substrates through TULIP2 methodology was not PCNA, which indicated that BRCA1/BARD1, and not RAD18, was the preferred ubiquitin E3 enzyme for PCNA under unperturbed conditions (Figure 2C). However, there was still the possibility that the absence of PCNA among the RAD18-specific substrates, regardless of the formation of the smear, was due to steric hindrance, similarly as previously observed for BRCA1 (Supplementary Figure 1).

To serve as positive control, we irradiated with UV light both BARD1-TULIP2 and RAD18-TULIP2 expressing Parental cells, performed the TULIP2 methodology and identified the substrates for both constructs after UV irradiation (Supplementary Dataset 4; Figure 2D). While PCNA enrichment as a BARD1-TULIP2 substrate was not affected by UV irradiation, in the case of RAD18-TULIP2, PCNA became now a substrate while it was not in unperturbed conditions, confirming the functionality of the RAD18-TULIP2 construct. Accordingly, PCNA ubiquitination levels were lower in BRCA1 knockout cells compared to parental cells in unperturbed conditions (Supplementary Figure 2B). But, after UV irradiation, ubiquitination kinetics of PCNA were not affected by BRCA1 knockout (Supplementary Figure 2C). Additionally, we confirmed by RAD18 knockdown, that PCNA ubiquitination after UV exposure was RAD18-dependent and BRCA1-independent (Supplementary Figure 2D). Finally, we performed the RAD18-TULIP2 assay in BRCA1-KO cells after UV irradiation (Supplementary Figure 2E). Mass spectrometry analysis revealed an additional enrichment of PCNA as a RAD18 substrate compared to parental cells (Supplementary Figure 2F).

Combined, these data allowed us to hypothesize that BRCA1/BARD1 and RAD18 share PCNA as substrate but act in different scenarios. While RAD18 would ubiquitinate PCNA when the replication fork encounters a replication fork barrier, BRCA1/BARD1 would perform PCNA ubiquitination at low levels to facilitate continuous DNA synthesis in unperturbed conditions (Figure 2E).

### BRCA1/BARD1 ubiquitinates PCNA to prevent ssDNA accumulation under unperturbed conditions

PCNA K164 ubiquitination has recently been shown to participate in replication fork protection (Thakar *et al*., 2020) and in preventing the accumulation of ssDNA gaps during DNA replication in unperturbed conditions (Leung et al., 2020), two functions that have been previously also attributed to BRCA1 (Billing et al., 2018; Cong et al., 2021; Daza-Martin et al., 2019; Panzarino et al., 2021). Different groups have shown that BRCA1-deficient cancer cells accumulate ssDNA gaps and thus can be therapeutically exploited with PARP or REV1 inhibitors (Cong *et al*., 2021; Taglialatela et al., 2021). We hypothesized that BRCA1/BARD1 mediates PCNA ubiquitination to protect replication forks and avoid the accumulation of ssDNA gaps in unperturbed conditions. Therefore, mutations on either PCNA or BARD1-BRCA1 that compromise its E3 activity could lead to ssDNA accumulation and genetic instability (Figure 3A).

**Figure 3.**
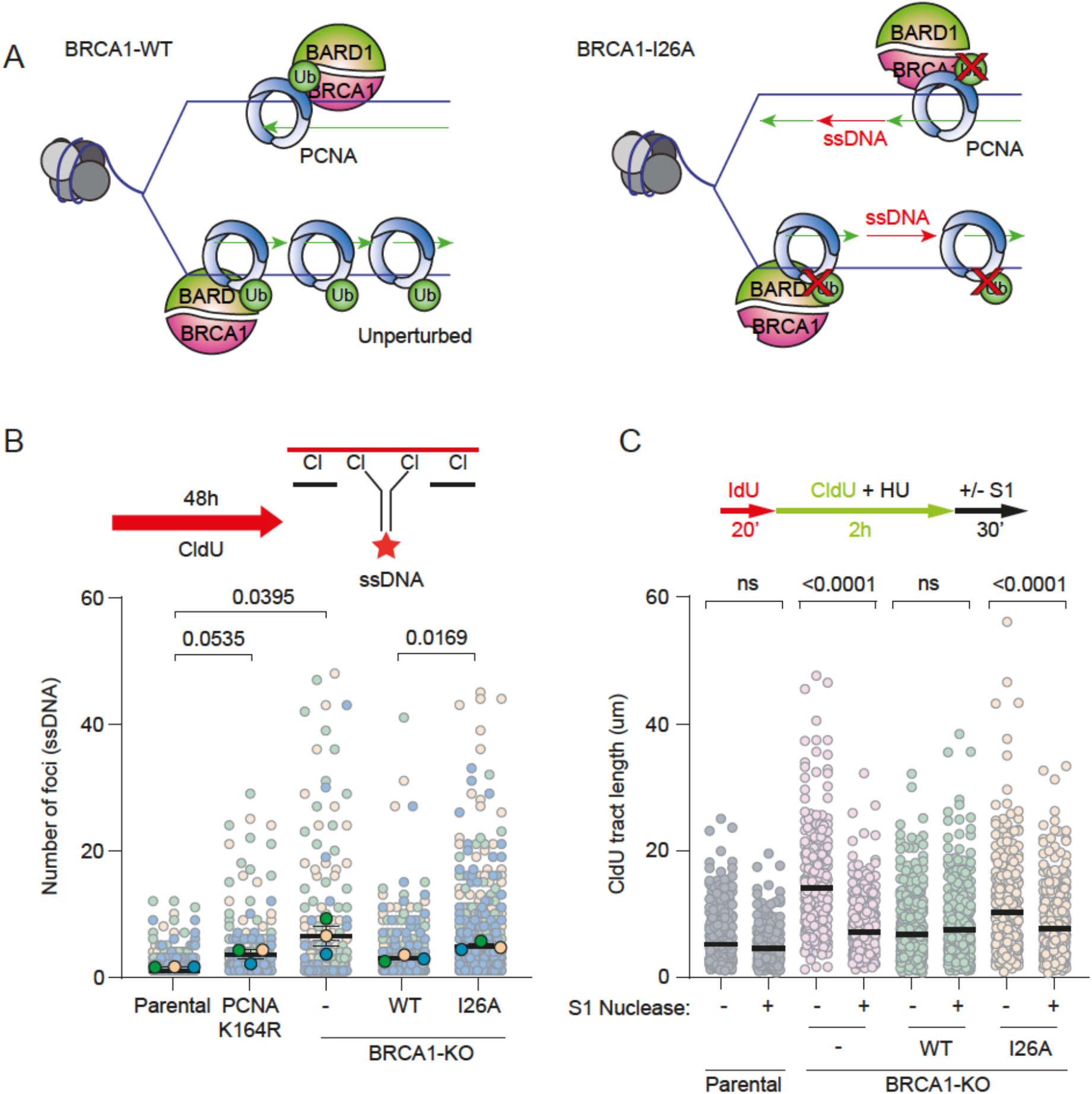
BRCA1-BARD1 mediated PCNA ubiquitination prevents ssDNA gaps. A. Proposed working model. While in wild type BRCA1 cells, BRCA1 would ubiquitinate PCNA in unperturbed conditions to facilitate continuous DNA synthesis, in an E3-deficient BRCA1-I26A mutant, deficiencies in PCNA ubiquitination in unperturbed conditions would promote the accumulation of ssDNA gaps. B. Cells were cultured in presence of CldU for 48h, fixed and analyzed by immunostaining against ssDNA. Number of ssDNA foci per cell is quantified. Each circle represents a cell. Average and standard deviation are depicted. Data corresponds to three independent experiments (N=3). p-values corresponds to two-tailed unpaired t-tests. C. Top: Scheme of the IdU/CldU pulse-labelling protocol, followed by S1 nuclease treatment. Bottom: CldU tract lengths in the indicated cell lines with and without S1 nuclease treatment. Each dot represents one fiber and the black bar represents the mean. At least 300 fibers were measured from two biological independent experiments (N=2). p-values were calculated using the Mann-Whitney test (ns: p>0.05).

To test BRCA1/BARD1 E3 activity on ssDNA gaps formation, we cultured different RPE1 cell lines with 10 μM CldU for 48 h and analyzed by immunofluorescence in non-denaturing conditions using an anti-BrU antibody that will only recognize CldU present in ssDNA gaps (Panzarino *et al*., 2021) (Figure 3B). In line with previous research, we observed an increased number of ssDNA gaps in BRCA1-KO and PCNA K164R mutant RPE1 cells under normal growth conditions (Cong *et al*., 2021; Thakar *et al*., 2020). BRCA1 cells reconstituted with BRCA1-GFP suppressed the formation of ssDNA gaps to the parental levels. However, complementation with the BRCA1-I26A-GFP mutant construct could not suppress ssDNA gaps formation and showed similar levels of ssDNA foci to PCNA-K164R mutant cells. Alternatively, to corroborate that the E3 activity of the BRCA1/BARD1 heterodimer prevents the formation of ssDNA gaps, we performed the S1-nuclease assay on DNA fibers (Figure 3C) in the presence of mild replication stress (Panzarino *et al*., 2021; Taglialatela *et al*., 2021). In this assay, DNA fibers containing ssDNA gaps would be shortened after treatment with S1 nuclease. Consistently, after mild replication stress (0.5 mM hydroxyurea for 2 hours), BRCA1-KO cells accumulated ssDNA gaps behind the replication fork, and this accumulation was suppressed after complementation with wild-type BRCA1-GFP but not by BRCA1-I26A-GFP.

### BRCA1/BARD1 E3 activity is required for replication fork protection and BRCA1/BARD1 isomerization

PCNA K164 ubiquitination protect stalled replication forks from nuclease activity and subsequent accumulation of ssDNA (Thakar *et al*., 2020). We hypothesized that upon replication fork blockade, deficiencies in PCNA ubiquitination by BRCA1/BARD1 will also result in fork de-protection and, due to nucleases activity, an increase in the exposed ssDNA, which can be measured by immunofluorescence. Therefore, we cultured Parental or BRCA1-KO cells complemented or not with either wild type BRCA1-GFP or BRCA1-I26A-GFP in the presence of CldU and treated them for 5 hours with 5 mM hydroxyurea to produce a complete replication fork blockade. Next, we analyzed the cells by immunofluorescence against RPA and exposed ssDNA (BrdU) and quantified the ssDNA signal (Figure 4A). Consistent with our hypothesis, BRCA1-deficient cells showed an increase in the intensity of ssDNA foci, and this effect was rescued by complementation with the wild type BRCA1-GFP construct but not with the BRCA1-I26A-GFP mutant construct.

**Figure 4.**
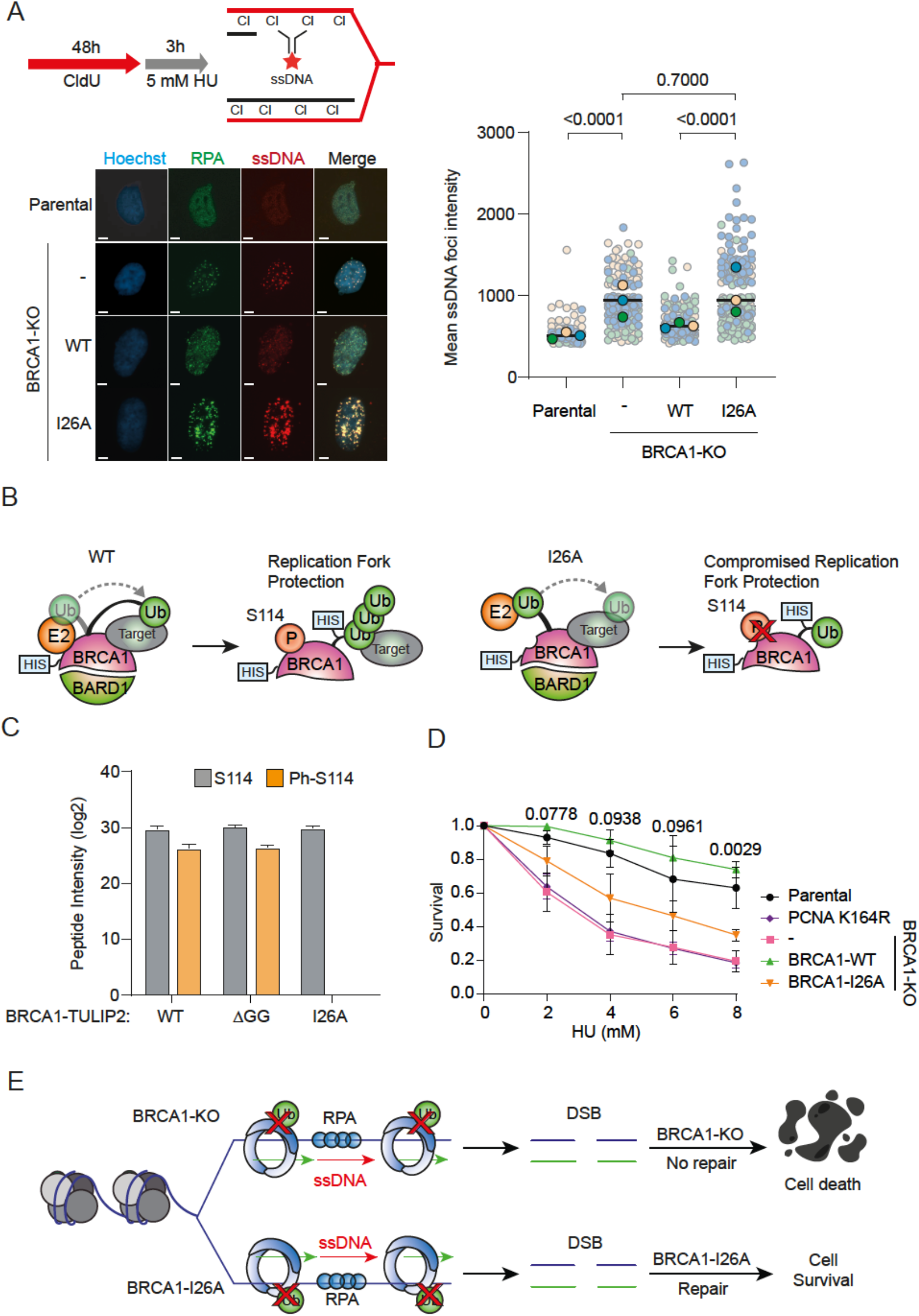
BRCA1/BARD1 E3 activity is required for replication fork protection and isomerization. A. ssDNA intensity in Parental and BRCA1-KO cell lines rescued with either BRCA1-WT or BRCA1-I26A. Cells were cultured in presence of CldU for 48 h and treated with 5 mM hydroxyurea for 3 h before fixation and immunofluorescence analysis against ssDNA. Size bars in fluorescence microscopy images represent 10 μm. Each point represents a cell. Three independent experiments were performed (N=3), average and standard deviation are shown. Average of each independent experiment is displayed with an orange, green or blue circle respectively for experiment 1, 2 and 3. p-values were calculated using the Mann-Whitney test. B. Schematic representation of S114 phosphorylation identified by MS in BRCA1-WT-TULIP2 and BRCA1-I26A-TULIP2. C. Intensities of tryptic peptide containing BRCA1 S114 in its unmodified or phosphorylated forms. Average of four independent experiments is displayed. Error bars correspond to Standard deviation. D. Survival assay of Parental, PCNA-K164R mutant and BRCA1KO cell lines rescued with either BRCA1-WT-GFP or BRCA1-I26A-GFP against different hydroxyurea concentrations. Three independent experiments with two technical repeats were performed (N=3). Standard deviation and p-values from unpaired t-tests comparing BRCA1-WT-GFP and BRCA1-I26A-GFP complemented cells are displayed. E. Model representing the relevance of BRCA1/BARD1 E3 activity in preventing ssDNA gap accumulation and promoting replication fork stability by PCNA ubiquitination. In the absence of BRCA1/BARD1 E3 activity, ssDNA gaps accumulate, which can eventually lead to the formation of DSBs. While in the absence of BRCA1, these DSBs would lead to cell death, in E3-deficient mutants, the DSBs can still be repaired by Homologous Recombination.

Replication fork protection upon replication stress by BRCA1/BARD1 requires BRCA1 phosphorylation at S114 which promotes isomerization of BRCA1/BARD1. BRCA1 S114 phosphorylation enhances the interaction of the BRCA1/BARD1 heterodimer with RAD51 and increases its loading at stalled forks, and consequently, BRCA1-S114 phosphorylation can be used as a reporter for BRCA1/BARD1 isomerization (Daza-Martin *et al*., 2019). BRCA1-S114A mutants are deficient in fork protection (Daza-Martin *et al*., 2019) similar to PCNA-K164R mutants (Thakar *et al*., 2020), and the BRCA1 ubiquitin E3-mutant I26A (Figure 4A). Therefore, we revisited the acquired mass spectrometry spectra of phosphorylation sites from our BRCA1-TULIP2 experiments (Supplementary Figure 1). In our experiment, we obtained good quality spectra for the tryptic peptide containing BRCA1-S114 both in its phosphorylated and non-phosphorylated form (Supplementary Figure 3A). However, while the signal for the unmodified peptide was similar for all three BRCA1-TULIP2 constructs, the I26A mutation prevented S114 phosphorylation from occurring (Figure 4B-C). This suggests that BRCA1 ubiquitin E3 activity is required for BRCA1 S114 phosphorylation and, as a consequence, BRCA1/BARD1 isomerization. Thus, we decided to test the sensitivity to hydroxyurea of the Parental and BRCA1-KO cells complemented or not with GFP-tagged constructs of either wild type or I26A mutant BRCA1. PCNA-K164R mutant was also included (Figure 4D). In contrast to what we observed for Olaparib (Figure 1A), where sensitivity of BRCA1-deficient cells could be rescued by complementation with the BRCA1-I26A mutant, in the case of hydroxyurea, while the complementation with the wild type GFP-BRCA1 construct could rescue the sensitivity of the BRCA1-deficient cells, the BRCA-I26A complemented cells were more sensitive than their wild-type counterpart. PCNA-K164R mutant cells were more sensitive than the I26A mutant, similar to BRCA1-deficient cells.

## DISCUSSION

Here, we provided novel insight in the role of the BRCA1/BARD1 ubiquitin E3 activity in the homologous recombination DNA repair pathway. Our results (Figure 1A-C) are in accordance with most studies and show that the E3 activity of BRCA1/BARD1 is not relevant for Homologous Recombination and Olaparib resistance (Nakamura *et al*., 2019; Reid *et al*., 2008; Shakya *et al*., 2011). However, we noticed that cells which were deficient in the BRCA1/BARD1 ubiquitin E3 activity were sensitive to hydroxyurea (Figure 4D) and accumulated more ssDNA in response to high concentrations of hydroxyurea that produce a total replication fork blockade (Figure 4A), which indicated that deficiencies in the E3 activity of BRCA1/BARD1 did not affect the DSB repair function of HR but do affect the replication fork protection function of HR (Tye et al., 2021).

Using TULIP2 methodology in unperturbed conditions, we identified PCNA-K164 as a ubiquitination substrate for BRCA1/BARD1 but not for RAD18 (Figure 1D-H; Figure 2C). Accordingly, previous studies show synthetic lethality in BRCA1 deficient cells with either RAD18 loss or the use of REV1 inhibitors (Nayak et al., 2020; Taglialatela *et al*., 2021). PCNA is known to be monoubiquitinated at low levels in mammalian cells in unperturbed conditions (Arakawa et al., 2006; Thakar *et al*., 2020; Unk et al., 2008), and PCNA ubiquitination levels in unperturbed conditions were reduced in BRCA1-deficient cells (Supplementary Figure 2B), although still some PCNA ubiquitination could be detected. E3s other than RAD18 and BRCA1 have been described, like for example Cullin 4 RING ligase, CRL4 (Terai et al., 2010). However, while CRL4 works synergistically with RAD18 and CRL4 knockdowns affect the accumulation of mono-ubiquitinated PCNA in response to UV irradiation (Terai *et al*., 2010), no differences were observed in PCNA ubiquitination in response to UV light in BRCA1-deficient cells (Supplementary Figure 2C-D), which supports the model that BRCA1/BARD1 and RAD18 modify PCNA in alternative pathways (Figure 2E).

Recently, it has been proposed that the synthetic lethality between BRCA1-KO and Olaparib treatments come from and increased replication fork speed in BRCA1 deficient cells and subsequent hyper accumulation on ssDNA gaps behind replication forks (Cong *et al*., 2021; Panzarino *et al*., 2021). The accumulation of ssDNA gaps behind the fork have been also detected in PCNA-K164R mutants (Thakar *et al*., 2020), and also when treating BRCA1-deficient cells with mild replicative stress like hydroxyurea concentrations not sufficient to produce a complete nucleotide depletion (Panzarino *et al*., 2021; Taglialatela *et al*., 2021). In this study, we show that the E3 activity BRCA1-I26A mutant show similar levels of ssDNA gaps formation as PCNA-K164R mutant and BRCA1-KO cells (Figure 3B-C), which seems, in principle, contradictory with the E3-deficient mutant being resistant to Olaparib treatment (Figure 1A; (Nakamura *et al*., 2019; Reid *et al*., 2008; Shakya *et al*., 2011)). We propose a model (Figure 4E) in which upon replication stress the BRCA1/BARD1 E3 activity would promote replication fork stability and continuous, ssDNA-free, DNA synthesis. Deficiencies in the E3 activity would eventually promote the appearance of DNA DSBs. While in BRCA1-deficient cells these DSBs would lead to cell death, in the E3-deficient mutants could still be repaired through homologous recombination-mediated DSB repair.

Noteworthy, the excessive replication fork speed observed in BRCA1-deficient mutants ((Cong *et al*., 2021; Panzarino *et al*., 2021; Taglialatela *et al*., 2021), Figure 3C), is, at least partially, rescued in the BRCA1-I26A mutant but not the ssDNA gaps formation. Thus, BRCA1/BARD1 inhibits the advance of replications fork in an E3-independent manner. We can speculate that this is mediated by the interaction of BRCA1 with the checkpoint protein ATR (Chiappa et al., 2022). Consistently, PARP inhibitors-resistant tumors, including BRCA1/2-revertant tumors which do not reconstitute RAD51 foci formation and Cyclin E-amplified tumors, are sensitive to ATR inhibition (Kim et al., 2020). Cyclin E overexpression also promotes ssDNA gap formation (Nayak *et al*., 2020). Accordingly, fork slowing is an adaptive mechanism to persistent ATR inhibition (Dibitetto et al., 2021).

Finally, some PARP inhibitor-sensitive BRCA1 tumors which mutations lead to disruption of the RING domain and a premature stop codon can revert by restoring the expression of a BRCA1 protein lacking the RING domain functionality but expressing the rest of the domains (Drost *et al*., 2011). Thus, treating these patients with other chemo therapeutic agents such as hydroxyurea, for which BRCA1/BARD1 E3-deficient cells are still sensitive (Figure 4D) might open a therapeutic opportunity for cancer treatment.

## ACKNOWLEDGMENTS

Authors thank Daniel Durocher and Sylvie Noordermeer for sharing the *TP53 ^-/-^* and *TP53^-/-^BRCA1^-/-^* hTERT RPE1 cell lines (Noordermeer *et al*., 2018) used in this work. This work was supported by a Young Investigator Grant from the Dutch Cancer Society (KWF-KIG 11367/2017-2) and the EMERGIA 2020 program (EMERGIA20_00276) from the Andalusian Regional Government-Junta de Andalucía, Spain to RG-P. Work in the laboratory of A.C.O.V. has been supported by the European Research Council (ERC; grant 310913) and the Dutch Research Council (NWO; grant 724.016.003).

## AUTHOR CONTRIBUTION

RG-P initiated the project. DS-L conducted most of the experimental work. LG assisted DS-L in RAD18-TULIP2 sample preparation. DS-L, LG and RG-P performed experiments in ACOV laboratory, NG-R performed the S1-nuclease fiber assay in PHS laboratory. AdR and PvV provided mass spectrometry data acquisition support. RG-P and AdR acquired mass spectrometry data. DS-L and RG-P analyzed mass spectrometry data. LG was supervised by DS-L, DS-L was supervised by RG-P. DSL and RG-P proposed and designed experiments and wrote the manuscript with input from other authors.

## METHODS

### Generation of TULIP2 constructs

BRCA1 and BARD1 genes were amplified and STOP codon removed by PCR using BP-tailed primers and cloned into pDNOR207 by Gateway® cloning BP reaction (Thermo Fisher Scientific). RAD18-pDNOR223 plasmid was obtained from Open Biosystems (Clone 1782) and the STOP codon was removed by site-directed mutagenesis. TULIP2 constructs were generated by Gateway® cloning LR reaction between donor plasmids containing BARD1, BRCA1 or RAD18 and acceptor TULIP2 plasmids (Salas-Lloret *et al*., 2019). Primers are listed in supplementary table 1.

### Cell culture

293T HEK and RPE1 cells were cultured in Dulbecco’s modified Eagle’s medium (DMEM) supplemented with 10 % Fetal Bovine Serum (FBS) and 100 U/mL penicillin/100 μg/mL streptomycin at 37°C and 5% CO2 unless specifically specified. Cells were regularly tested for mycoplasma contamination.

### TULIP2 Lentivirus production

293T HEK cells were seeded at 30% confluency in a T175 flask containing 17 mL of DMEM + 10% FBS. After 24 hours, the transfection mixture was prepared by combining lentiviral packaging plasmids 7.5 μg pMD2.G (#12259, Addgene), 11.4 μg pMDLg-RRE (#12251, Addgene), 5.4 μg pRSV-REV (#12253, Addgene) and 13.7 μg TULIP2 plasmid with 114 μL of 1 mg/mL Polyethylenimine (PEI) in 2mL 150 mM NaCl. The mixture was vortexed and incubated 15 min at room temperature before adding to the HEK cells. Next day, culture medium was refreshed with DMEM 10% FBS. 72 hours post-transfection, lentiviral suspension was harvested by filtering through a 0.45 μm syringe filter (PN4184, Pall Corporation) and kept at −20 °C for further use. Lentiviral particle concentration was determined using the HIV Type 1 p24 antigen ELISA Kit (ZeptoMetrix Corporation).

### Generation of TULIP2 and stable 10xHis-Ub RPE1 cell lines

RPE1-hTERT *TP53^-/-^* and RPE1-hTERT *TP53^-/-^ BRCA1^-/-^* cells were kindly provided by Dr. Sylvie Noordermeer (Verver et al., 2016) and were seeded in 15 cm diameter plates at 10% confluency with DMEM 10% FBS. Next day, cell culture medium was replaced with lentiviral TULIP2 constructs containing medium with 8 μg/mL polybrene.

After 24 hours, medium was refreshed with DMEM 10% FBS, 1% Pen/Strep. 72 hours post-infection, TULIP2 positive clones were selected on puromycin.

RPE1-hTERT *TP53^-/-^* and RPE1-hTERT *TP53^-/-^ BRCA1^-/-^* cells were infected using a bi-cistronic lentivirus encoding 10xHis-Ub and Puromycin resistant gene separated by an IRES (Cuijpers et al., 2017) and selected on puromycin after 72 h.

### Purification of TULIP2 conjugates

Following the TULIP2 methodology (Salas-Lloret *et al*., 2019), five 15 cm diameter plates of RPE1 cells containing a TULIP2 construct, were grown up to 60% to 80% confluence. Expression of TULIP2 constructs was induced with 1μg/mL doxycycline once 60-80% confluence was reached. 24 h after doxycycline induction, cells were non-treated (NTC) or treated with different treatments. Proteasome inhibitor MG132 (Sigma Aldrich) was used at 10 μM for 5h, olaparib (Bio-connect) and bleomycin (Millipore) for 2 h at 10 μM and 5 μg/mL respectively. Treatment was performed at 40 J/m^2^ UV with 5 hours recovery.

After a particular treatment, cells were washed twice with ice-cold PBS and scraped. Next, cells were spun down and collected in 5 mL ice-cold PBS, 100 μL of sample was taken as input and lysed in 200 μL SNTBS buffer (2% SDS, 1% NP-40, 50mM TRIS pH 7.5, 150 mM NaCl). After additional centrifugation, cells were lysed in 10 mL Guanidinium buffer (6M guanidine-HCl, 0.1M Sodium Phosphate, 10mM TRIS, pH 7.8) and snap frozen in liquid nitrogen. After thawing, lysates were homogenized at room temperature by sonication at 80% amplitude during 5 s using a tip sonicator (Q125 Sonicator, QSonica, Newtown, USA). Sonication was performed twice. Subsequently, protein concentration was determined by BiCinchoninic Acid (BCA) Protein Assay Reagent (Thermo Scientific). After equalization, lysates were supplemented with 5 mM β-mercaptoethanol and 50 mM Imidazole pH 8.0. 100 μL of dry nickel-nitrilotriacetic acid-agarose (Ni-NTA) beads (QIAGEN), were equilibrated with Guanidinium buffer supplemented with 5 mM β-mercaptoethanol and 50 mM Imidazole pH 8.0. Equilibrated Ni-NTA beads were added to the cell lysates and incubated overnight at 4°C under rotation.

After lysate-beads incubation, Ni-NTA beads were transferred with Wash Buffer 1 (6 M Guanidine-HCl, 0.1M Sodium Phosphate, 10 mM Tris, 10 mM Imidazole, 5 mM β-mercaptoethanol, 0.2 % Triton X-100, pH 7.8) to an Eppendorf LoBind tube (Eppendorf). Subsequently, beads were washed with Wash buffer 2 (8 M Urea, 0.1M Sodium Phosphate, 10 mM Tris, 10 mM imidazole, 5mM β-mercaptoethanol, pH 8) and transferred to a new LoBind tube with Wash buffer 3 (8 M urea, 0.1M Sodium Phosphate, 10 mM TRIS, 10 mM imidazole, 5 mM β-mercaptoethanol, pH 6.3). Ultimately, beads were washed twice with Wash buffer 4 (8 M urea, 0.1M Sodium Phosphate, 10 mM TRIS, 5 mM β-mercaptoethanol, pH 6.3). After last wash, Ni-NTA beads were resuspended in 100 μL of 7 M urea, 0.1 M NaH2PO4/Na2HPO4, 0.01 M Tris/HCl, pH 7 and 10% of the sample was taken as pull down for immunoblotting.

### Lys-C and trypsin digestion of TULIP2-purified conjugates

Ni-NTA beads were firstly digested with 500 ng recombinant Lys-C (Promega) at RT while shaking at 1,400 rpm. After 5h with Lys-C, urea buffer was diluted to <2M by adding 50 mM ABC. A second digestion was performed o/n at 37°C while shaking at 1,400 rpm using 500 ng of sequencing grade modified trypsin (Promega). Trypsin digested peptides were separated from Ni-NTA beads by filtering through a 0.45 μm filter Ultrafree-MC-HV spin column (Merck-Millipore).

### Mass Spectrometry sample preparation

Digested peptides were acidified by adding 2% TriFlourAcetic (TFA) acid. Subsequently, peptides were desalted and concentrated on triple-disc C18 Stage-tips as previously described (Rappsilber et al., 2007). Stage-tips were in-house assembled using 200 μL micro pipet tips and C18 matrix (Sigma Aldrich). Stage-tips were activated by passing through 100 μL of methanol. Subsequently 100 μL of Buffer B (80% acetonitrile, 0.1% formic acid), 100 μL of Buffer A (0.1% formic acid), the acidified peptide sample, and two times 100 μL Buffer A were passed through the Stage-tip. Elution was performed twice with 25 μL of Elution buffer (32,5% acetonitrile, 0.1% formic acid solution).

Samples were vacuum dried using a SpeedVac RC10.10 (Jouan, France) and stored at −20°C. Prior to mass spectrometry analysis, samples were reconstituted in 10 μL 0.1% formic acid and transferred to autoload vials.

### LC-MS/MS data acquisition

Mass spectrometry data was acquired either by and nanoLC Easy 1000 (Proxeon, Odense, Germany) coupled to a Q-Exactive mass spectrometer (Thermo, Bremen, Germany) (BARD1-TULIP2 samples) or Ultimate 3000 nano-gradient HPLC system (Thermo, Bremen, Germany), coupled to an Exploris480 mass spectrometer (Thermo, Bremen, Germany) (BRCA1-TULIP2; RAD18-TULIP2 and BARD1-TULIP2 with UV irradiation).

For the Q-Exactive, chromatography was performed as in (Salas-Lloret *et al*., 2019), peptides were separated in an in-house packed with Reprosil-Pur C18-AQ 1.9 μm (Dr. Maisch, Ammerbuch, Germany) 20 cm analytical column in a 45 minutes gradient from 0% to 30% acetonitrile gradient in 0.1% Formic Acid followed of 20 minutes of column re-equilibration. The mass spectrometer was operated in a Data-Dependent Acquisition (DDA) mode with a top-7 method and a scan range of 300–1,600 m/z. Fullscan MS spectra were acquired at a target value of 3 × 10^6^ and a resolution of 70,000, and the Higher-Collisional Dissociation (HCD) tandem mass spectra (MS/MS) were recorded at a target value of 1 × 10^5^ and with a resolution of 35,000, an isolation window of 2.2 m/z, and a normalized collision energy (NCE) of 25%. The minimum AGC target was 1 ×10^4^. The maximum MS1 and MS2 injection times were 250 and 120 ms, respectively. The precursor ion masses of scanned ions were dynamically excluded (DE) from MS/MS analysis for 20 s. Ions with charge 1, and >6, were excluded from triggering MS2 analysis.

For the Exploris480, samples were injected as in (Salas-Lloret et al., 2022) onto a cartridge precolumn (300 μm × 5 mm, C18 PepMap, 5 μm, 100 A) with a flow of 10 μl/min for 3 minutes (Thermo, Bremen, Germany) and eluted via a homemade analytical nano-HPLC column (50 cm × 75 μm; Reprosil-Pur C18-AQ 1.9 μm, 120 A) (Dr. Maisch, Ammerbuch, Germany). The gradient was run from 2% to 38% solvent B (80% acetonitrile, 0.1% formic acid) in 120 min. The nano-HPLC column was drawn to a tip of ~10 μm and acted as the electrospray needle of the MS source. The temperature of the nano-HPLC column was set to 50°C (Sonation GmbH, Biberach, Germany). The mass spectrometer was operated in data-dependent MS/MS mode for a cycle time of 3 seconds, with a HCD collision energy at 28 V and recording of the MS2 spectrum in the orbitrap, with a quadrupole isolation width of 1.2 Da. In the master scan (MS1) the resolution was 120,000, the scan range 350-1600, at an standard AGC target with maximum fill time of 50 ms. A lock mass correction on the background ion m/z=445.12 was used. Precursors were dynamically excluded after n=1 with an exclusion duration of 45 s, and with a precursor range of 10 ppm. Charge states 2-5 were included. For MS2 the scan range mode was set to automated, and the MS2 scan resolution was 30,000 at a normalized AGC target of 100% with a maximum fill time of 60 ms.

### Mass Spectrometry data analysis

All raw data was analyzed using MaxQuant (version 1.6.7.0) as previously described (Tyanova et al., 2016a). Search was performed against an *in-silico* digested UniProt reference proteome for Homo sapiens including canonical and isoform sequences (24th January 2022). Database searches were performed according to standard settings with the following modifications: Digestion with Trypsin/P was used, allowing 4 missed cleavages. Oxidation (M), acetyl (protein N-term), phospho (S,T) and GlyGly (K) for ubiquitination sites were allowed as variable modifications with a maximum number of 3. Label-Free Quantification (LFQ) was enabled, not allowing Fast LFQ while permitting iBAQ and matching between runs.

Output from MaxQuant Data were exported and processed for statistical analysis in the Perseus computational platform version 1.6.7.0 (Tyanova et al., 2016b). LFQ intensity values were log2 transformed and potential contaminant and proteins either identify by site or only reverse peptides were removed. Samples were grouped in experimental categories and proteins not identified in 4 out of 4 replicates in at least one group were removed. Missing values were imputed using normally distributed values with 0.3 width and 1.8 down shift separately for each column. After imputation, statistical analysis was performed using two-sided Student’s t tests. Results were exported into in MS Excel for a comprehensive browsing and visualization of the datasets. Volcano plots were constructed for data visualization using the VolcaNoseR web app (Goedhart and Luijsterburg, 2020) (https://huygens.science.uva.nl/VolcaNoseR2/).

### LC-MS/MS data availability

The mass spectrometry proteomics data have been deposited to the ProteomeXchange Consortium via the PRIDE (Perez-Riverol et al., 2019)partner repository with the dataset identifier PXD039167

### Purification of 10xHis-Ubiquitin conjugates

10xHis-Ub conjugates were purified using Ni-NTA beads as previously described (Hendriks and Vertegaal, 2016). In brief, RPE1 cells were grown in three 15 cm dishes and were scraped and lysed in 6M guanidinium buffer. A small fraction of cells was separately lysed in SNTBS buffer as input control. After homogenization by sonication, lysates were incubated with Ni-NTA beads o/n at 4 °C. Next day, Ni-NTA beads were washed with buffers 1-4 and eluted in 7 M urea, 0.1 M NaH2PO4/Na2HPO4, 0.01 M Tris/HCl, pH 7.0, 500 mM imidazole pH7.

### Electrophoresis and immunoblotting

Samples were separated on Novex 4-12% gradient gels (Thermo Fisher Scientific) using NuPAGE® MOPS SDS running buffer (50mM MOPS, 50mM Tris-base, 0.1% SDS, 1mM EDTA pH 7.7) and transferred onto Amersham Protran Premium 0.45 NC Nitrocellulose blotting membranes (GE Healthcare) using a Bolt Mini-Gel system (Thermo Fisher Scientific), which was used for both the gel electrophoresis and the protein transfer to the membrane according to vendor instructions. Membranes were stained with Ponceau-S (Sigma Aldrich) to determine total amount of protein loaded. Next, membranes were blocked with blocking solution (8% Elk milk, 0.1% Tween-20 in PBS) for 1 h prior to primary antibody incubation. Chemiluminescence reaction was initiated with Western Bright Quantum Western blotting detection kit (Advansta-Isogen) and measured in a ChemiDocTM imaging system (BIO-RAD, Hercules, CA, USA). Antibodies are listed in supplementary table 2.

### Clonogenic survival assay

RPE1 cells lines were seeded at 3000 cells/well in 6-well plates and allowed to attach overnight. TULIP2 constructs were induced with 1 μg/mL doxycycline prior treatment. Olaparib (Bio-connect) was added at different concentrations for 24 hours. After treatment, medium was refreshed. Hydroxyurea (Sigma) was added at different concentrations and medium was refreshed after 16 hours treatment. Subsequently, cells were allowed to grow for 10 days and fixed for 20 minutes in 4% paraformaldehyde (PFA) in PBS. Cells were stained with Crystal Violet 0.05% for 30 minutes and washed with water. Afterward, Crystal Violet was re-solubilized in methanol and O.D.595 was measured in the VICTOR X3 Multilabel Plate Reader 2030-0030 (Perkin Elmer). GraphPad was used for statistical analysis and the value of untreated cells was set at 100% survival.

### Immunofluorescence

RPE1 cell lines were seeded at 20% confluency on 8 mm coverslips in 12-well plates and allowed to attach overnight. 1 μg/mL doxycycline was added to required cell lines to induce TULIP2 constructs 24 hours prior treatment. For RAD51 foci experiments, cells were treated with 10 Gy with 4 h recovery before fixation. For single-stranded DNA (ssDNA) gaps, medium was supplemented with 10 μM CldU (Sigma). After 48 hours, medium was replaced with either 10 μM olaparib or 5 mM hydroxyurea and cells were fixed with 1% PFA, 0.3% Triton X-100, 0.5% methanol after 3 hours treatment. RAD51 experiments were performed using a Leica SP8 confocal microscope taking 8 frames per image. ssDNA gaps experiments were performed in a ZEISS fluorescent microscope. ImageJ and GraphPad were used for quantification and statistical analysis.

### Quantification and statistical analysis

Quantification of microscopy data was performed using Fiji-ImageJ and the statistical analysis was performed in GraphPad Prism 8. Statistical details of individual experiments can be found in figure legends, including the statistical test performed and definition of center and dispersion representation. For every analysis, N represents the number of values considered in the statistical analysis.

### S1 nuclease assay

Cells were pulse-labeled with 20 μM IdU (20 min), washed twice with PBS and pulse-labeled with 200 μM CldU in the presence of 0.5 mM HU for 2h. Cells were then washed twice with PBS and permeabilized with CSK buffer (100 mM NaCl, 10 mM MOPS pH 7, 3 mM MgCl2, 300 mM Sucrose and 0.5% Triton X-100 in water) for 8 min at RT. Permeabilized cells were treated with S1 nuclease buffer (30 mM Sodium acetate pH 4.6, 10 mM Zinc acetate, 5% glycerol, 50 mM NaCl in water) with or without 20 U/ml S1 nuclease (Invitrogen, 18001-016) for 30 min at 37°C. Cells were then scrapped in PBS + 0.1% BSA, pelleted and resuspended in PBS + 0.1% BSA at a final concentration of 1-2×10^3^ cells/μl. 2.5 μl of cell suspension were spotted on a positively charged slide and lysed with 7.5 μl of spreading buffer (200 mM Tris-HCl pH 7.5, 50 mM EDTA, 0.5% SDS). After 8 min, slides were tilted at 45 degrees to allow the DNA to spread. Slides were then air-dried, fixed with ice-cold methanol/acetic acid (3:1) for 5 mins, air-dried and stored at 4°C. Slides were rehydrated with PBS, denatured with 2.5 M HCl for 1h, washed with PBS twice, and blocked with blocking buffer (3% BSA, 0.1% Triton X-100 in PBS) for 40 min. Next, slides were incubated with primary antibody mix of mouse anti-BrdU which recognizes IdU (Becton Dickinson #347580, 1:250) and rat anti-BrdU which recognizes CldU (Abcam #6326, 1:250) diluted in blocking buffer for 2.5h at RT in a dark humid chamber. Slides were washed 3 times with PBS for 5 min each and incubated with secondary antibodies anti-mouse Alexa fluor 594 and anti-rat Alexa fluor 488 (1:250, Invitrogen #A11005 and #A11006, respectively) in blocking buffer for 1h at RT in a dark humid chamber. After washing 3 times with PBS and air-drying, slides were mounted with Prolong gold antifade reagent (Invitrogen, P36930) and stored at 4°C until imaging. Images were acquired using a AF6000 Leica Fluorescence microscope equipped with a HCX PL APO 63x (NA =1.4) oil objective. At least 300 fibers per condition were measured using the segmented line tool on ImageJ FIJI software (https://fiji.sc).

