## Supplemental Figures for "BRCA1/BARD1 ubiquitinates PCNA in unperturbed conditions to promote replication fork stability and continuous DNA synthesis"

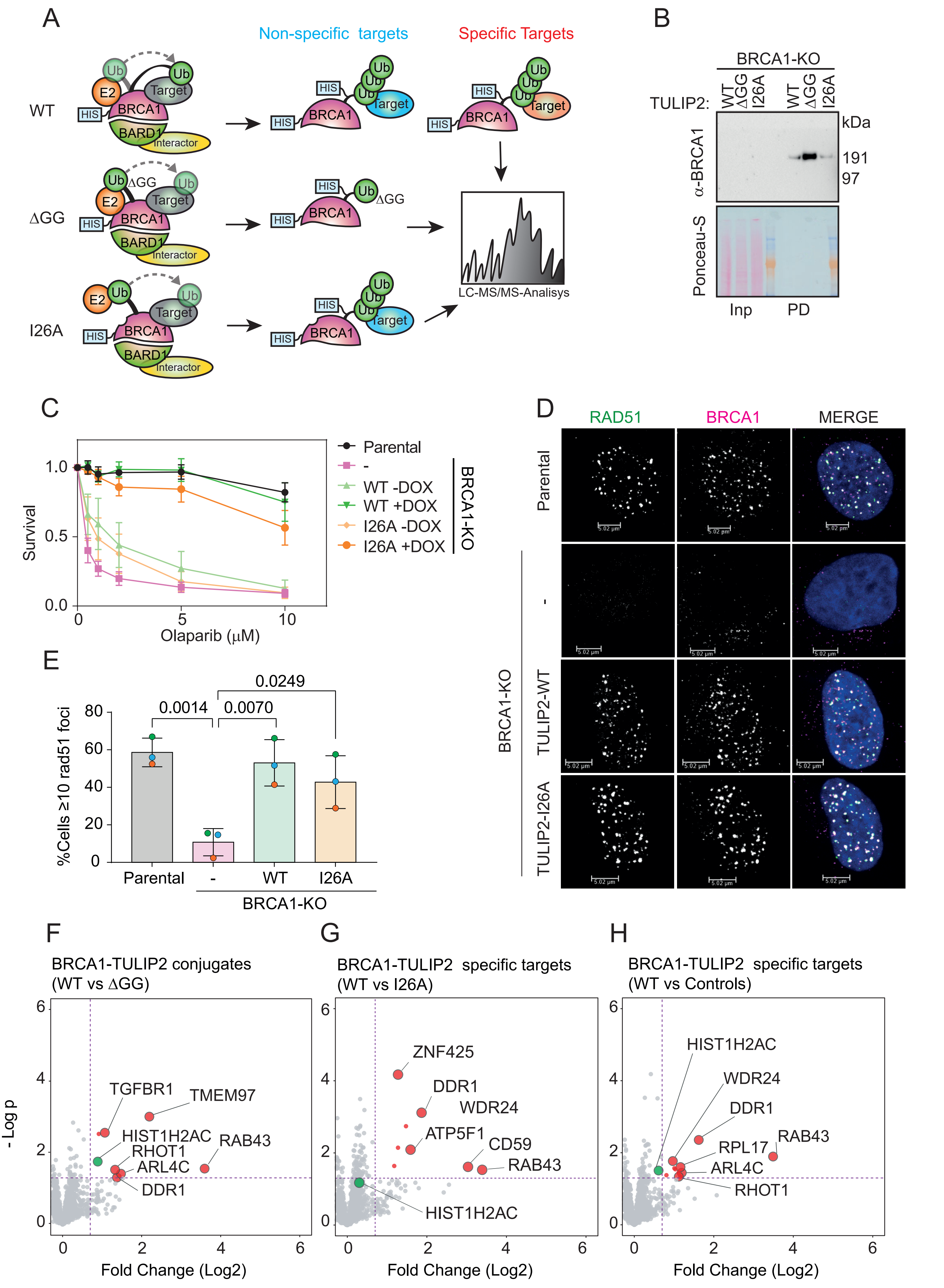


**Supplementary Figure 1**

**Figure S1**. **BRCA1-TULIP2 Characterization**. **A**. BRCA1-TULIP2 Rationale to find BRCA1 specific targets using Mass Spectrometry. **B**. Analysis by immunoblotting of TULIP2 pull downs in BRCA1-KO cells rescue with BRCA1-TULIP2 constructs (WT, ∆GG and I26A). **C**. Survival assay after treatment with Olaparib of Parental and BRCA1-KO cells resued or not with either WT or I26A mutant BRCA1-TULIP2 constructs. Four independent experiments with 5 technical repeats were performed (N=4). Average and standard deviations are displayed. **D, E**. Analysis by immunofluorescence against RAD51 and BRCA1 of Parental and BRCA1-KO cells rescued or not with either WT or I26A BRCA1-TULIP2 constructs. Quantification of percentage (%) of cells with equal or more than 10 RAD51 foci (D) and representative images (E) are provided. Size bars in fluorescence microscopy images represent 10 µm. Three independent experiments were performed per condition displaying the average and standard deviation (N=3). The average of each independent experiment is represented by an orange, green or purple circle. P-values correspond to two-tailed unpaired t-tests. **F**. Volcano plot depicting statistical differences between BRCA1-WT and ∆GG TULIP2 constructs. Each dot represents a protein **G**. Volcano plot depicting statistical difference between BRCA1-WT and BRCA1-I26A TULIP2 constructs. Each point represents a protein. **H**. Volcano plot depicting statistical difference between BRCA1-WT-TULIP2 samples compared to ∆GG and BRCA1-I26A TULIP2 samples as controls. Each dot represents a protein.

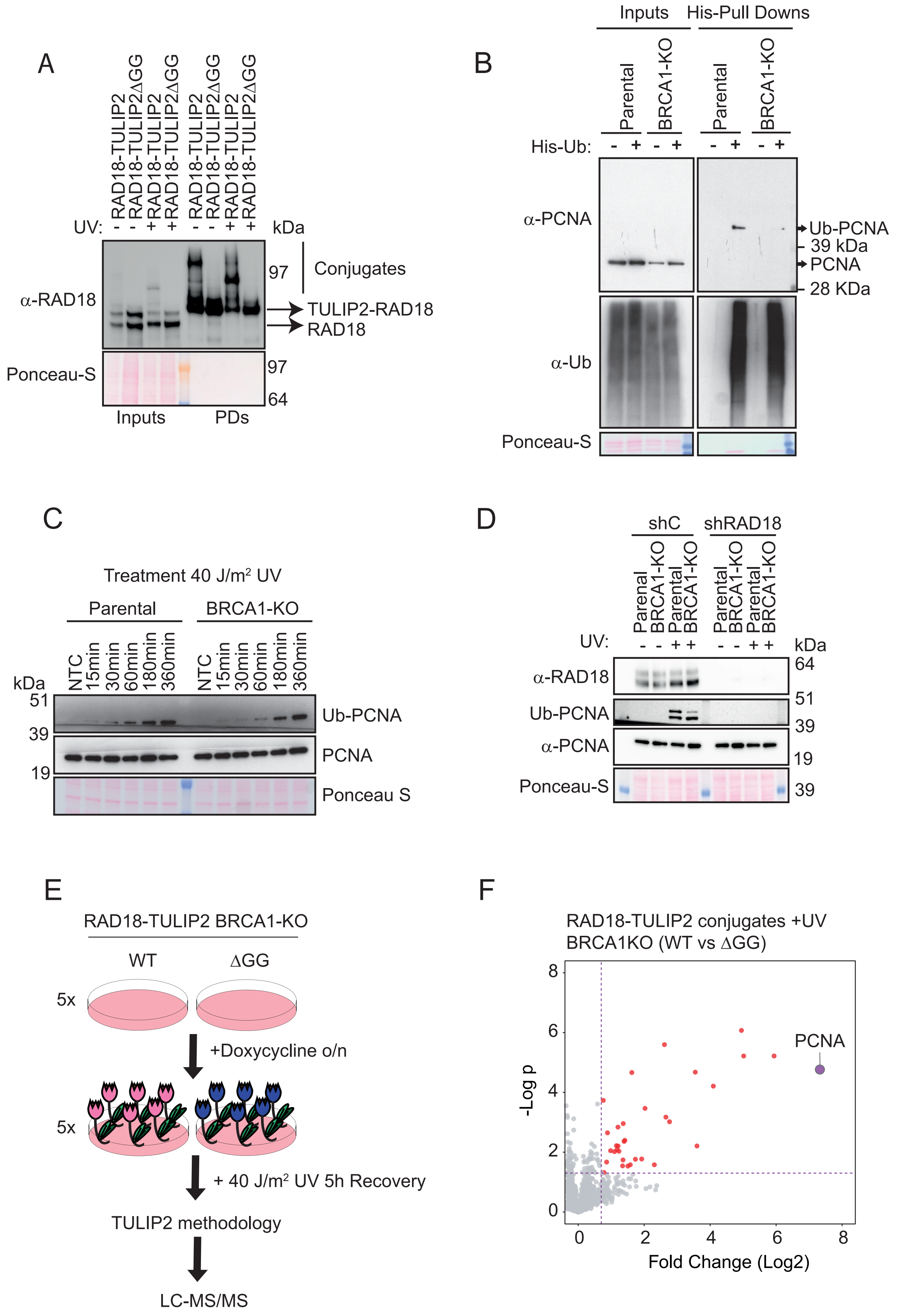


**Supplementary Figure 2**

**Figure S2**. **(A)** Analysis by immunoblotting of RAD18-TULIP2 samples with and without 40 J/m$^2$ UV treatment. **(B)** Analysis by immunoblotting of the HIS-Ubiquitin proteome in Parental and BRCA1-KO cells. **(C)** Analysis by immunoblotting of PCNA ubiquitination after 40 J/m$^2$ UV irradiation in a time course manner in Parental and BRCA1-KO cells. **(D)** Analysis by immunoblotting against PCNA 5h after 40 J/m$^2$ UV irradiation in Parental and BRCA1-KO cell after treating with a control or RAD18-targetting shRNA. **(E)** Experimental setup for RAD18-TULIP2 methodology in BRCA1KO cells 5h after 40 J/m$^2$ UV irradiation. **(F)**. Volcano plot depicting statistical differences between RAD18-TULIP2 samples and RAD18-∆GG-TULIP2 samples in BRCA1KO cells 5h after 40 J/m$^2$ UV irradiation. Each dot represents a protein, PCNA is highlighted.

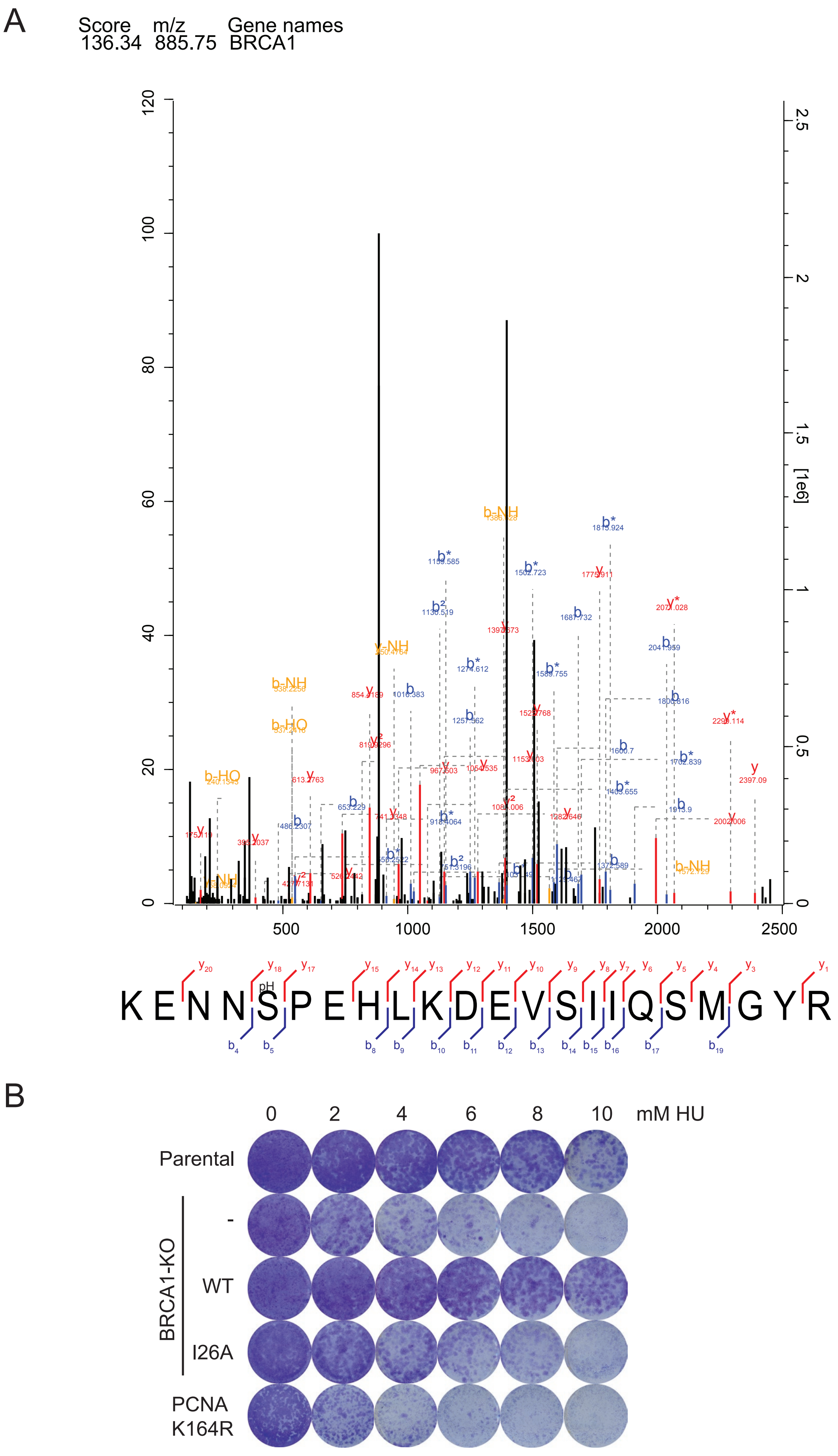


**Supplementary Figure 3. A.** Acquired spectrum corresponduing to the best localized peptide for BRCA1-S114 phosphorylation. **B**. Survival assay of Parental, PCNA-K164R mutant and BRCA1-KO cell lines recued with either BRCA1-WT or BRCA1-I26A against different hydroxyurea concentrations. Three independent experiments with two technical repeats were performed (N=3). A representative repeat is displayed.
